# Neuroecological selection for musical features through spatial reciprocity in long-term partnerships

**DOI:** 10.1101/2022.12.13.520337

**Authors:** David M. Schruth

## Abstract

Chemical sensing via olfaction is an ancient form of inter-organism communication. And acoustical signaling via tonal and rhythmic patterning is common among vertebrates. Those animals that live in diffused habitats and move in diasporic ways have further evolved more spectrally varied and discretized calls. But unlike song in birds, researchers have struggled to locate isolated nuclei specialized for cognition for music in humans. The brain stem, midbrain, hindbrain, and forebrain, however, all largely associate with aspects of musical performance, perception, memory, and emotion. I hypothesized that spectral features of musical display evolved as honest signals of spatial cognition for precarious locomotor tasks associated with nurturing and protecting vulnerable offspring. I investigated possible connections between motor, visual, and spatial cognitive areas in relation to both signaler production and receiver processing of acoustical features of musical output. Brain component volume fractions of 42 parts from 48 primates were compiled, from a single source, and compared against a vocal complexity index as well as individual musical feature scores: tone, interval, repetition, transposition rhythm, and unique syllable count. Structures for spatial and visual perception, as well as motor control and emotional processing, associated moderately with areas used by species who produce calls with both temporal and spectral musical features. These findings are consistent with a dual (both receiver- and signaler- side) function of musical signals. Associations with spatio-social areas support direct selection for a paralimbic-based neighbor-orienting sensory modality, possibly for mapping and anticipating movement of fellow arboreal cohabitants. Associations with motor areas support the complementary model that signaler capacities for spatio-motor emplacement are indirectly selected by conspecific receivers. This dual manifestation in low-parity species that locomote in diasporic ways through diffuse habitats, is compatible with musicality serving as courtship signals by long-term mates with consistent and reliable spatial capacities relevant to care of [arboreally] vulnerable offspring.

## Introduction

Vertebrate production of acoustic vibrations audible across great distances is realized via the passing of air over constricted nasal or throat passages (Stevens, 2000). These, often tonal, waveforms are regularly used to communicate position and emotive state to conspecifics, other niche-inhabitants, or predators. Added variation, as in interval height, can aid in signals being heard in more occluded, arboreal environments (Morton, 1975). Other modificaitons to sound signals, such as spectral diversity in vocal units, can help to communicate a unique vocal signature of identity to conspecific receivers (Owren, Seyfarth and Cheney, 1997).

At the even more elementary, neural-cellular level, foundations for both vision (in the form of rods and cones) and hearing (in the form of cochlea [c] hair cells) are known to be stable and extremely ancient (500mya) facets of vertebrate biology (Jerison, 2000). The production and perception of tone also likely pre-dates the emergence of mammals. Even some aspects of rhythm (beat entrainment) are thought to be nearly 300 million years old (Honing, 2018). And, notably, monotonic rhythm routinely appears in calls of most non-human primate species (Schruth and Jordania, 2020).

More complex, music-like patterning, approaching acoustic display, might also possibly function as a signal (Mehr *et al*., 2020) perhaps of underlying cognition. Clustering of repeated or transposed motifs requires receivers (and learners) of such calls to possess cognition for matching of acoustic patterns (Schulz *et al*., 2008). High-level synthesis of acoustic signals occurs in auditory cortical regions of the temporal lobe (Morley, 2012). The hippocampus, basal ganglia, and cerebellum also house circuitry utilized for fusing of sound gestures containing melodic features (Levitin, 2006; Jordain 1997; Fernández, 2015). But acoustic stimulii are typically first received in the cochlea, and are then projected, via the spiral ganglion and nerve c. to the c. nucleus of the brain stem (Sando, 1965).

Poorly understood, still, is how these animals’ brains are used to process musical features (Jacoby *et al*., 2020). Birds, who are the most well known for song, are endowed with specialized nuclei for singing (Moore *et al*., 2011). Primates also have proportionally large brains that correspond to long, slow (and perhaps learning conducive) life-histories (Jones, 2011). Tarsiers, like birds, have exceedingly small cerebrum, and may be less dynamic in their vocal capacities than other primates (Nietsch and Niemitz, 1987). No special music or song nuclei exist in primates and their rather stereotyped calls tend not to be highly influenced by social learning (Mitani, 1988). Deducing the distributed set of brain structures consistently involved in musical calling has therefore proven challenging.

### Brain Areas

Cognitive scientists have spent decades piecing together how sounds that reach the ear trigger the cascade of mechanical, chemical, and neural events in various brain regions to produce auditory perception (Peretz and Zatorre, 2005). The production of musical sounds has a likewise complicated cascade of neural projections but in the opposite direction, from neural to mechanical. Following my predecessors, I segregated these various processes into the four principal functional domains of production, perception, memory, and emotion. Within each category, I highlight the brain regions shown to be active corresponding to these functional domains across numerous studies and sources. A key study that served as an important starting point was for musical processing of pleasurable as well as displeasing sounds (Blood and Zatorre, 2001).

### Production

The music-like sounds of animals are generated just like many other vocalized sounds, by constricting air passages (e.g. larynx, vocal folds, syrinx) to create vibrations corresponding to different wavelengths, waveforms, and acoustic patterning. Preceding this mechanical sound production, motor-neural impulses project from the pons by virtue of vocal pattern generators. These projections pass through the cerebellum in the hindbrain, as well as the peri-aquiductal gray [PAG] and tegmentum in the midbrain. and thalamus (Peretz and Zatorre, 2005). Additional studies have shown varying degrees of association with subcortical nuclei of, and surrounding, the basal ganglia (Ackermann, Hage and Ziegler, 2014), including the substantia nigra, striatum, and meso-cortical cingulate. In the neocortex, both the frontal and sensory motor cortex have been implicated. While studies on production of specific musical features may be more elusive and inconsistent, it is evident that nearly all levels of the brain are involved in creating musical sounds, possibly also across disparate taxa.

### Perception

Periodic concentrations of air molecules reach the ear as sound patterns and are funneled towards the tympanic membrane. These patterns of air molecules induce vibrations which may reach optimal resonance with particular bands of hair cells corresponding to particular frequencies of the initiating sounds. Frequency information is individually relayed from the cochlear spiral ganglion and is projected onto the cochlear nerve towards the cochlear nucleus of the brain-stem. From here, nerve impulses make a u-turn back up the brain-stem towards higher brain structures including the olive near the cerebellum, colliculus in the brain-stem, and the sub-cortical thalamus. At any point along the way, reflexive short-circuiting may bypass higher brain oversight by reacting in a way that prioritizes time-sensiive survival-conducive responses to potentially dangerous stimuli.

Although both spectral and temporal features of sound are essentially (rhythmically) periodic, they are typically processed in different areas of the higher brain. The temporal lobes play a crucial part of processing spectral information. Harmonic and pitch information has been associated with processing by Heschl’s gyrus in the superior part of the temporal lobe. The posterior secondary cortex has been implicated in melodic analysis and processing pitch changes. Melodic processing and recognition of contours and pitch intervals are all facilitated by both the left and right temporal lobes.

Intermediate to the lower brain relaying of auditory signals and higher brain integrative processing of harmonic, intervallic, and melodic information, the mid-brain performs a basic spatial mapping of sound sources in two dimensions. Archicortical areas such as the hippocampus and surrounding periallocortical areas of the entorhinal cortex include the schizocortex as well as the pre- and para-subiculum. These areas typically facilitate orientation and navigational functionality, although lower (mid-brain) areas such as the tectum and higher cortical areas such as the insula, also facilitate this process of mapping sensory input to times and places. The insula was, until recently, considered to be just another part of the temporal lobe, while others have interpreted its extreme medial position to be essentially paralimbic. Having six cortical layers, it is now considered a fifth lobe in its own right, though it is deeply folded inside the lateral sulcus, underneath three other lobes, including the temporal, parietal, and frontal as well as the orbital operculum. The insula is the least well understood of all of the other neocortical lobes, but *is* understood to be a specialized integrator of disparate multi-modal sensory information. This sub-temporal lobe integrates such varied external sensations into a cohesive gestaltic picture of the surrounding (socially relevant) environment, but still within the context of proprioceptive internal states. Thus, the insula is ideally situated to act as a multimodal and interpersonal organ of socio-ecological decision making.

### Memory

Like the more medial paralimbic areas accessory to the hippocampus (including the schizo- and entorhinal-cortex), the insula serves as a memory hub for temporal, orientational, and navigational capacities. Unlike these lower cortical areas, the insula specializes in connecting space-time coordinates with other more socioemotional and sensorimotor relevant information—attached, for example, to places and faces. In addition to self-monitoring physiological needs such as thirst, hunger, pain, and fatigue, the insula is responsible for governing socioemotional capacities for compassion, empathy, and interpersonal experience. Lastly, the insula plays an important role in higher order self-awareness and risky decision-making. Musical memory typically implies memory for form and structure of events along with minimal contextual, rather than emotional, associations. But I introduce the full function of the insula and integrate with these other areas in the next section.

Temporal-adjacent structures such as the insula and Heshl’s gyrus are primary cognitive regions involved in musical perception and memory (Peretz and Zatorre, 2005). Other neocortical regions, however, such as the temporal and frontal lobes, are also involved in such faculties. Dorsolateral and inferior frontal areas are most often recruited areas for when working memory load is high. Memory is critical not only for connecting stimulus with context, but also for higher order matching and sound stream splitting and pattern recognition. Birds and humans have converged upon abilities to perform rhythmic and perceptual grouping (ten Cate and Spierings, 2019) which underlie learning and participation in complex sonic environmental contexts. In larger groups, where musical displays are longer and more rhythmic, there are typically a larger number of contributors (as opposed to smaller familial contexts) in which musical memory becomes all the more critical.

Cerebral storage space constraints also factor into accommodating the open-ended process of longer songs and larger song repertoires that tend to arise through the process of sexual selection (Slater, 2000). Thus, memory also plays an important role in social (inter-sexual) signaling by vertebrates, a topic which receives more attention below.

### Emotions

Auditory stimuli are maintained in working memory in order to compare an element in a sequence to one that occurs later in time. Emotional processing of musical experience is also neurally isolable. Expression of tone, mode, and tempo are manifestations of musical emotion that correspond to particular brain areas. That is, emotional analyses of musical stimuli are mediated through a spectral-perceptual cortical relay. Nearly the entire brain has nuclei involved in emotional associations with music. The dorsal midbrain, hindbrain cerebellum, limbic hypothalamus and amygdala, subcortical nucleus accumbens and ventral striatum, as well as orbito-frontal, insular, and (pre) cuneus of the occipital lobes all process emotional aspects of music. Theorietical efforts have attempted to delineate the more emotionally melodic from the more lexically rhythmic (Peretz & Colthart 2003). Moving anteriorly from the more physio-mechanical parts of the temporal lobe, toward more socioemotional parts near the frontal lobe, Broca’s area, a hub for spoken language, may be implicated (Brown, Martinez and Parsons, 2004).

The study of relationships between musical behavior and emotion are as old as the earliest theories on the subject. Theorists predating the ideas of natural and sexual selection proposed that music was merely an advanced form of physical expression of emotion—a manifestation of “nervous excitement” (Spencer, 1875). Since the resurgence of ideas in the origins of music, many others have joined onto this line of thought. Modern ideas on the function of musicality place it alongside other artistic behaviors as essential to ritual ceremonies that relieve emotional anxiety and coordinate members of a group (Dissanayake, 2009), managing social uncertainty, facilitating affiliative and pro-social interaction (Cross, 2011), or coordinating affect for social identity, cooperation and alliance building (Bryant, 2013). Others have reframed music as vocal and kinesthetic communication of socioemotional content (Morley, 2014), as increasing value through emotional contagion, social cohesion, improved well being (Snowdon, Zimmermann and Altenmüller, 2015), or as serving to alleviate stress by reducing the costs of group living (Savage *et al*., 2021).

### Hormones

Less work has been done to map hormonal correlates of musicality. Many studies have looked at the relationship between testosterone, the male sex hormone, and music performance in people ranging from non-musicians to elite musicians. Recently some have looked in more detail at how hormones interface with particular regions of the brain. For example, testosterone is thought to positively stimulate activity in both the amygdala and hypothalamus (Miani, 2015), whereas oxytocin, the love (or bonding) hormone, is thought to stimulate activity in both the hippocampus and orbitofrontal cortex. This line of evidence has been used to support the idea of musicality as evolving as an ornament for courtship (Miani, 2015).

### Spatial modules for motor control and conspecific orientation

The four functional domains can be matched to two mechanisms of evolution. The first is for performance related to motor control of the vocal apparatus. The second relates to perception, memory, and emotion (but also learning) of such acoustic output. I have recently proposed names to accompany this sender-receiver pair of signaling modules: as motive emplacement and acoustic neighbor orientation.

### ME

This module of “motive emplacement” refers to the motor control required to realize arbitrary targets in space, but also to the motor control required to create the acoustic signals that convey such an ability. Thus motive emplacement is not only a reference to locomotor and positional targeting of substrate and dietary items, but the vocal manifestation of motor control required by small muscles to achieve such skeletal motor tasks. Further, since musical animals have eyes that are constantly re-oriented via rectus muscles, accurate function of such fine motor control (hidden behind cranial bones), may be signaled by equally small (but more publicly showcased) muscles of the vocal tract. Thus the skill in quickly maneuvering and focusing an essential (yet invisible) organ for feeding, moving, and survival, could be signaled by another motoric organ that may more readily be assessed, assuming that overlapping (mid-) brain (-stem) circuitry governs both.

### PIANO

A second receiver-side module is also proposed and is hypothesized to reside in the medial temporal lobe as both paralimbic and insular. This sub-cortical system, surrounding and including parts of the limbic system (e.g. the hippocampal gyrus), primarily implicates the *piriform*, *parahippocampal*, and *entorhinal* cortices. These ancient structures overlap with the lowest three layers of the temporal lobe, *hippocampus*, and the olfactory structures of the *piriform*. While the *piriform cortex* is primarily concerned with sense of smell, the other two, along with the adjacent *hippocampus*, are important in facilitating spatiotemporal navigation as well as memory encoding and retrieval. Interestingly, the *schizocortex*—containing the presubiculum, parasubiculum, (involved in directing of head position and used in spatial navigation), and the enthorhinal cortex—also shares contiguity with these other spatial orientating neural areas. This proposed module also includes the aforementioned insula, an understudied fifth lobe of the neocortex. The insula is the ideal candidate for a higher-cortical target of gestaltic processing of perceptual stimuli: integrating visual, olfactory, and auditory information into a coherent whole. This ultimate projection may be more socioemotionally sensitive to affiliation than the lower areas (e.g. amygdala) that are more proximate to a fight-or-flight brain-stem level response. Analogous reactions may also occur for threatening chemical stimulii entering the fornix or for low-frequency sounds entering the cochlea. Higher anterior cingulate cortex and the adjacent septum have terminal targets associated with maternal bonding and emotional, motivational, and goal-directed behavior. Thus, the final reaches of the anterior insula might be similarly inhibitively egalitarian and socioemotionally aware.

### Evolutionary Mechanisms

#### Human musicality and various selection models

Theories about the origins of music can be partitioned into two recently productive schools of thought. One posits it has evolved as a by-product of individualistic survival, while the other sees it as adaptive (or exaptive) in a more pro-social capacity.

For example, music could be seen as manipulating our evolved perception for “attention worthy events” (Pinker, 1997). An example is “shrieks” and “screams” that can trigger a startle response (Bryant, 2013). We do know that dissonant sounds can easily scare us, while consonant sounds have an attractive affect (Fritz *et al*., 2009). Indeed, tonal calls tend to correspond to states of high arousal and may prescribe reorienting towards sources of sounds. Thus, the prime mover of diversity in human musicality could merely relate to manipulation of an ancient need to navigate the sonic environment. Our ancestors met this need by segregating sound streams, locating sound sources, and categorizing sounds with efficiency and accuracy (Bregman, 1990; Bryant, 2013).

On the opposite extreme, theories of a social bonding purpose are prevalent. Bonding ideas—ranging from those within pairs, groups, mothers and infants— abound in the literature. One of the primary arguments supporting bonding hypotheses suggests that the repeat cost of duet-learning with a new partner or group is an honest and costly signal of partner investment that inherently discourages disillusionment (Geissmann and Orgeldinger, 2000). In all bonding theories, an ultimate objective tends to be for the group signal to act as advertisement of the bond strength to external groups (Geissmann and Orgeldinger, 2000).

An alternative category for origins ideas rests upon the stable foundation of signaling theory. Numerous signal types are invoked under these forms of evolutionary argument. Signals can be costly, or honest, or both. They can be indicators of an underlying trait, possibly indicating general vigor or motor control. A new version of this form of signaling applies to signaling between groups in relation to musical display, and has been dubbed “credible signaling.”

Within evolutionary ideas, there are also numerous forms of selection ranging from natural (on individuals) or selection acting on pairs or groups. Selection can also be asymmetrical, as in the case of sexual selection, with one sex, typically females, choosing the other, typically males. Roles can also be more egalitarian, as in mate-choice or “sexual choice” (Miller, 2001). Under sexual selection, secondary (sexual) traits will tend to become exaggerated especially if there are large asymmetries in parental investment (Trivers, 1972) and males have little to offer females beyond their genes.

Evolution by (usually female) mate choice can be direct, indirect, or non-selective. Under direct selection, females choose males who can provide territory or parental care—something I have termed “nurtural security”—whereby food sources can be readily converted into offspring. Indirect selection merely has females selecting conceptions by males for purely genetic reasons. Under non-selection, males tend to exploit biased sensitivities merely for copulation opportunities and songs tend to develop novel features in order to avoid habituation by potential mates.

#### Bird song signaling and selection for motor control

A number of studies of birds have confirmed a connection between many of the above performance features of song and some combination of earlier pairing and higher (extra-pair) copulations or fertilizations (Searcy and Yasukawa, 2019).

Partial evidence exists for non-selective, bias-exploitative, or “ornamental” signaling, driving singing in birds, which persists as a prevalent mechanism as it tends to increase copulation opportunities. On the other hand, there may be stronger evidence for motor control via indirect signaling for “conditional” indicators of viable offspring outcomes from mating (Searcy and Yasukawa, 2019). In primates these might be motor control, for landing limbs, or for understanding parabolic trajectories for leaps (Schruth, 2022).

In birds, motor performance tends to reflect (indirect) genetic skill or vigor [SoV], whereas ornamental traits arise secondarily as a way to enhance this SoV (Byers 2010). Examples include repertoire size, syllable diversity, or other performance indicators: time, length, rate, and amplitude (Gil and Gahr, 2002) as well as unit repetition or trill interval consistency. These vocal displays are often also accompanied by wing, leg, or locomotor displays. Subtle differences in such displays can serve as useful indicators of developmental history, genetic viability, and current health or condition (Byers, Hebets and Podos, 2010).

#### Bird song signaling and mate choice

Given that singing and musicality in birds and primates (respectively) tends to occur in monogamous species, I chose to explore various mate-choice related evolutionary mechanisms (Searcy and Yasukawa, 2019). Direct selection mechanisms for mate choice have somewhat less evidence in birds. Examples include species recognition or decreased cost in energy or risk of choosing a mate. Direct mechanisms can also include honest handicaps (e.g. features of acoustic signals) that likely mediate both conspecific recognition and mate selection. (Wilkins 2013) These direct signals translate to viable offspring via both dietary factors such as provisioning, territory defense, or other forms of care, even if the chosen partner doesn’t contribute genes (Searcy and Yasukawa, 2019). Conveniently, in primates, egalitarian monogamous mating systems are common correlates of musicality (Schruth, Templeton and Holman, 2021). For mutual benefit, direct selection mechanisms should manifest through reciprocity between partners (esp. monogamous pairs) for energetic pay off in slowly reproducing species with low parity birthing. Honesty of “revealing” or costly handicap indicators is maintained by a finite cognitive storage capacity (Gil and Gahr, 2002), which can also be highly spatial in modality.

#### Spatial reciprocity and mate choice in gibbons

A model example of such spatial reciprocity exists in gibbons. These socially monogamous primates hang precariously from branches, using just a single hand at a time, as a part of brachiating, their primary form of locomotion. Gibbons also pair with mates for life, thus making selecting a mate an especially important choice that can take many years to cohere. Similarly, since they experience concentrated weight per digit (Congdon and Ravosa, 2016) as well as acceleration during landing, such forces are further compounded when gestating or carrying infants. While there is some day-to-day trade-off in infant carrying duties, females bear the brunt of the burden, especially in the first few months of the infant’s life. Males instead may act to offset this energetic burden via defense of the vulnerable mother and infant, as well as scouting food sources and boundaries of their ever-shifting home range. This reciprocity of spatial responsibilities might include reliable motive emplacement and leap landing, as well as effective memory and orientation abilities for accurately localizing a new mate and neighbors for within-tree and range spacing logistics. Long-term partnerships in complex and precarious settings require both long-term memory and motor planning (and accurate execution) that can readily access corresponding higher-brain memory. Additionally, these brain structures should be further along a given neural pathway from other more reflexive, lower-brain, fight-or-flight functionality.

#### Hypotheses: primate musicality via sexual choice and mate selection

I hypothesized that musicality evolved as both indirect and direct signals involved in mate selection for long-term partnerships. First, locomotion between canopy elements requires reflexive spatiomotor execution in target landing tasks given the precarious gravitational constraints of narrow inter-branch targets and precipitous above-ground distances. I predicted a bias of auditors selecting for conspecifics who produce consistently reappearing units in display calls, thereby indirectly selecting for genes that encode quality motor-control capacities also recruited for rapid grasp landing. Receivers of such signals, like the call-learners themselves, should possess perceptual cognition for accurately comparing and grouping these acoustic units. Second, longer-term pairings between inter-substrate arboreal cohabitants likely requires individuals to possess fully functioning spatial, proprioceptive, orientational, navigational, and turn-taking capacities. Given the [k-]selective advantages of predation avoidance and the excessive g-forces involved in (esp. infant-laden) landing of such inter-branch locomotion, periods of weaning and learning by infants should be protracted. Thus, regardless of siring certainty, long-term mate selection should directly entail choosiness for cortical function in spatially reciprocating partners capable of role flexibility and payoff postponement in nurturing and protecting [arboreally carried] offspring (Table 2).

**Table 1.**
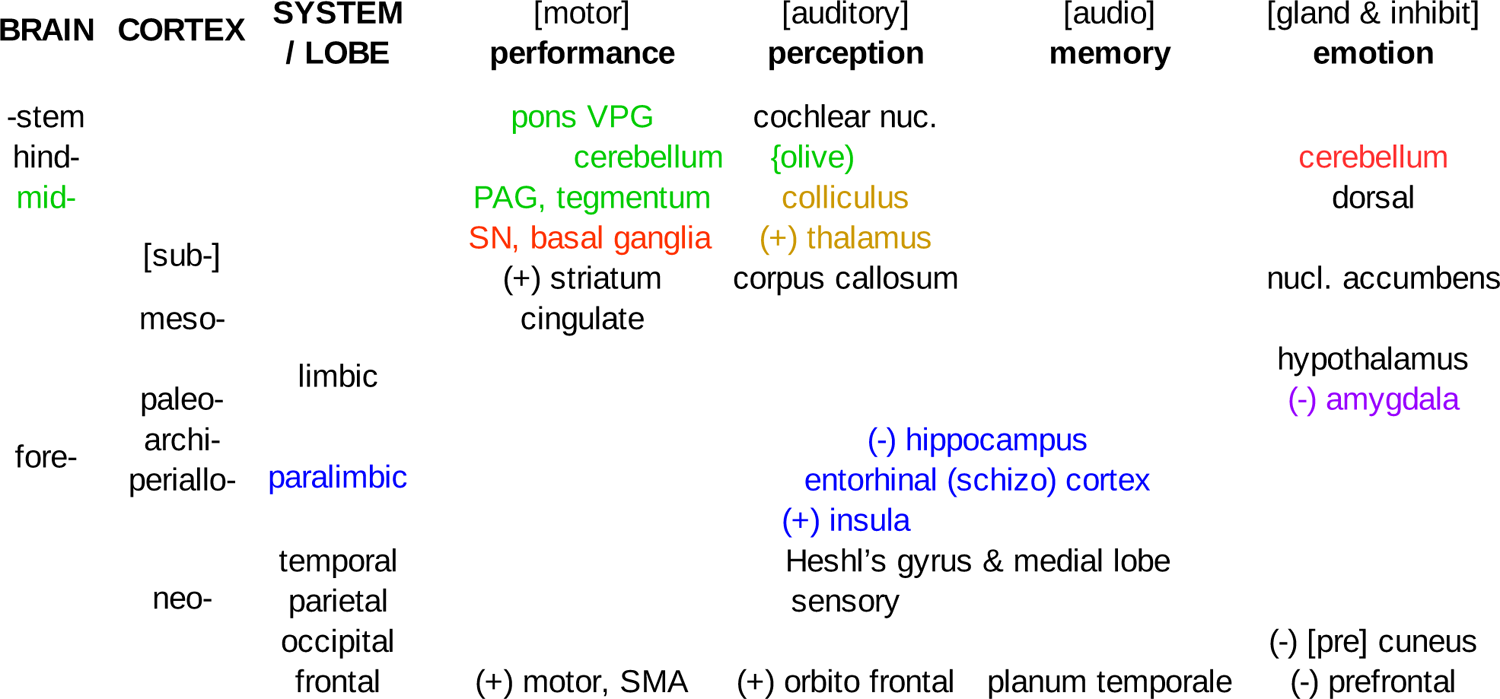
Brain areas involved in musical processing by functional domain For each brain level, cortex and system-specific cortical structures involved in musical processing are implicated in relation to the four functional domains of performance, perception, memory, and emotion. Colored text corresponds to a number of proposed modules as presented in this text: blue corresponds to the PIANO module components whereas brown, green, and red correspond to motive emplacement [ME] modules. Red specifically corresponds to rhythmic components. Areas prefixed with a plus (or minus) symbol correspond to positive (or negative) activation in response to consonant (or dissonant) sounding musical stimuli, according to various studies by Zatorre.

**Table 2.** Mate choice, selection, signaling, and processing associations. Sexual selection can be partitioned into three sub-types corresponding to direct, indirect, (or even non-selective) payoffs to mating decisions. While indirect indicators of motive emplacement [ME] as performance signals, for example, may be more difficult to measure, they have the most theoretical support in explaining the evolution of musical calling. Direct, reciprocal signals of [mostly future] parental care are thought to be perceived by the PIANO module. Non-selective by-products of ornamental signals can also evolve via increasing chances of copulation by exploiting auditory perception centers that are biased toward emotionally sensitive response.

| selection type | signal | payoff | (long-term) benefit | processing | functionality |
| --- | --- | --- | --- | --- | --- |
| direct | reciprocal | energetic | ♀ parental care & ♂ nurtural protection | perception | P.I.A.N.O. |
| indirect | indicator | genetic | ♂ motor & ♀ parabolic cognition | performance | M.E. & L.L. |
| non-selection | ornamental | copulation | variability ↔ anti-habituation | emotional | M.E.E. & A.C. |

#### Music, musicality, and feature component evolution

Music is a complex whole-brain phenomenon created primarily by humans. Darwin suggested it was the most mysterious of all our behaviors (Darwin, 1871). Others have suggested that, because its function has an indeterminate [adaptive] role as a whole, we should look instead for the mechanistic function of each incremental addition of individual components throughout its historical evolution (Honing *et al*., 2015). For example tone/pitch, beat perception, and metrical rhythm likely evolved one at a time over the course of (hundreds of) millions of years. This multi-component and pluralistic approach naturally also lends itself to a comparative analytical methodology while also employing an ecological perspective for selection on traits (Fitch, 2015). Elsewhere, I have developed an acoustic reappearance diversity index as such a continuous measure of musicality which can be easily used in comparative analyses across species. ARDI retains relevance to minimally capturing the complex whole of musical behavior in humans, but is less useful, other than as a starting point, in addressing the multi-component aggregation of individual musical features. While I predicted that features of reappearance and diversity are the most theoretically relevant, in testing ideas connecting musicality with ecologically relevant spatial abilities for long term partnerships, rhythm and tone may also be important. Thus this paper also investigates all six (collected) musical features as independent factors individually consociating with ecological and neurological aspects of musicality.

## Materials and Methods

I calculated brain component volume fractions on a small sample of non-human primates. Volumetric data on 42 different brain parts from 48 primate species was complied from a single source (Matano et al., 1981). Part fractions were calculated by dividing the volume of each part by the overall brain volume. These were compared to a vocal complexity index and six structural acoustical features prevalent in human music and bird song. The diversity of reappearing (unique) units [ARDI] was measured as the probability of syllable reappearance—calculated as repetition plus transposition times the syllable count itself—and is used synonymously here as a measure of musicality (Schruth, Templeton and Holman, 2021). Ordinary least-squares linear regression was used to assess associations of brain volume fractions and their aggregates with both the individual musical feature scores and ARDI (Figs. 1 & 3). Regressions against ARDI were also performed with and without *Tarsier spp.* (Fig. 3, colored lines), as this species exhibited several signs of being an outlier, albeit an important one for inferring ancient evolution. Overlap in these datasets ranged from *n*=22 to *n*=7, variably depending on brain sub-component availability. For this reason, analysis was also performed at the general brain functional level, combining several brain components, under a weighted average, to generate composite cognition outcome variables. Functional brain regions were categorized and weighted (decreasing monotonically) according to the lists in Table 4.

**Figure 1.**
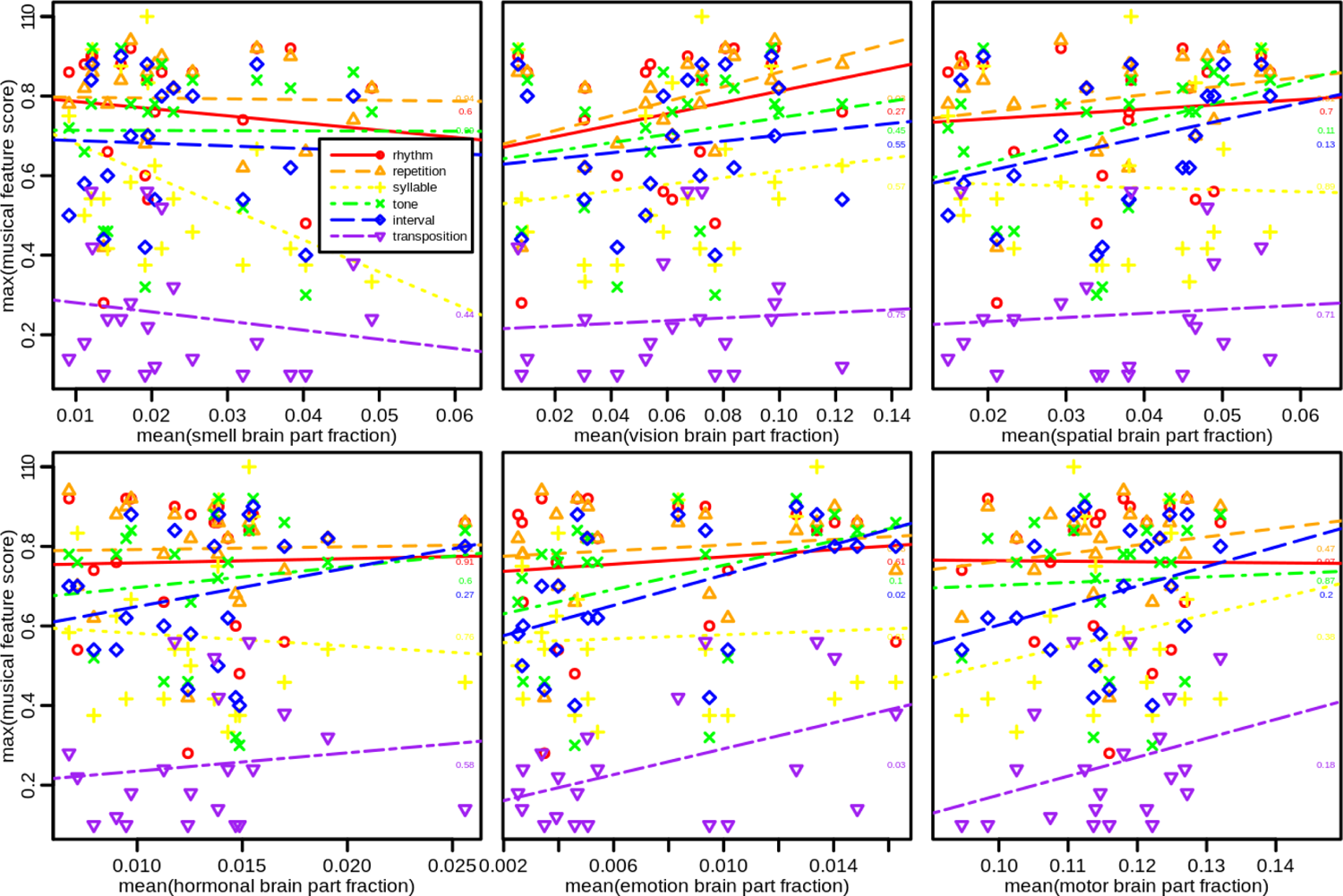
Weighted mean brain function versus max musical feature scores for six brain functions. Using the functional brain area definitions (Table 4) monotonically decreasing weighted means of these fractions were compared with the maximum musical feature scores for each species. *Syllable count* was rescaled to 0 to 1 to facilitate comparison with the other features. Linear regression fit lines were plotted across each of the six feature score datasets.

**Table 3.** Glossary of acronyms and terms

|  |  |
| --- | --- |
| **A.R.D.I.** | acoustic reappearance diversity index {continuous index of manifest musicality measuring the average number of repeated syllable types in a call} |
| **M.E.** | motive emplacement {setting and attaining arbitrary targeted goals in space} (2021) |
| **M.E.E.** | motive emplacement and entrainment {intentionally realizing body movements with respect to socially generated acoustical cues and signals} (2022) |
| **L.L.** | leap landing: a locomotor form of motive emplacement whereby leaps are landed on narrow arboreal substrate targets |
| **A.A.H.** | acoustic adaptation hypothesis {features evolved to improve propagation of acoustic signals around obstacles specific to particular habitats} (Morton 1975) |
| **A.S.A.** | auditory scene analysis {perceiver-side recovery of individual sound streams from a cacophonous mixture of multiple environmental sounds} (Bregman 1994) |
| **A.C.** | “auditory cheesecake” (Pinker 1994) or auditory chocolate, whereby biased sensitivities for environmental listening is exploited by easily processed musical sounds |
| **P.I.A.N.O.** | paralimbic insular acoustic neighbor orientation {higher-cortical para-hippocampal brain areas involved in orientation and navigation tasks that facilitate spatial harmony with conspecifics and cohabitants of shared ecological niches} (2021 & 2022) |
| **k-selection** | selection in species characterized by slow (body & population) growth rates, low parity, long life spans, long weaning durations, and relatively altricial offspring |
| **sexual selection** | competition between members of the same sex (detering rivals) in order to attract mates (Darwin 1871); can result in bizarre and “ornamental” secondary traits (Miller 2001) |
| **sexual choice** | more sexually-symmetrical form of mate choice via fitness indicators, in contrast to more asymmetrical sexual selection based on exaggerated or costly signals (Miller 2001) |
| **mate choice** | mate selection broadly construed around the full life-cycle: including courtship, copulation, fertilization, gestation, and parenting (Westneat & Fox 2010) |
| **mate bonding** | a possibly acoustic mode of acquaintance before cohabiting (e.g. a tree) or parenting; a part of courtship and pair formation (adapted from Savage 2020) |
| **pair bonding** | long term bonds formed by established couples typically jointly rearing offspring, post copulation; thought to co-evolve with vocal duetting (Wickler, 1980) |
| **mate selection** | process of choosing long-term mates in k-selected species (includes most of the mate choice cycle except offspring nurturing—the signal payoff) |
| **signal selection** | Includes selection for costly / honest signals (sent to all rivals, partners, enemies); but excludes utilitarian selection (e.g. fighting ability) (Zahavi & Zahavi 1997) |
| **nurtural security** | providing for offspring via both menage and range defense as well as dietary scouting or provisioning (can include infant carrying) |
| **trophic security** | evolutionary or developmental means (e.g. infant carrying) of successful deterrence or avoidance of predation within a given food web (Schruth & Jordania 2020) |
| **spatial reciprocity** | long-term mutual benefit between breeding partners or long-term neighbors where spatial-boundaries are respected in order to avoid physical collisions and falls (e.g. within tree) or resource shortages and conflicts (e.g. between trees) |

**Table 4.** Functional brain areas and their underlying components

|  |  |
| --- | --- |
| **smell** | bulbus olfactorius, lobus piriformis, bulbus olfactorius<br>accessorius, nucleus olfactorii lateralis |
| **vision** | area striata, visual cortex, tractus opticus, corpus geniculatum<br>laterale, nucl_amygd_basal_IMC, vestibularis medialis |
| **spatial** | hippocampus, schizocortex, septalis triangularis,<br>nucl_amygd_basal_IMC, subcommissurale |
| **motor** | mesencephalon, thalamus, basal ganglia ,cerebellum,<br>nucl_amygd_basal_IMC |
| **emotion** | amygdala, pineale, septum, habenularis medialis, complexus<br>cortico basolateralis |
| **hormonal** | hypothalamus, pineale, septum, meninges hypophysis nerves |

## Results

There were negative associations of vocal complexity with brain components associated with olfaction—including the *bulb*, *accessorius*, and *piriform*—and was strongest between *piriform* and *syllable* (*p*<0.05). Inversely, there were uniformly positive associations of all musical features with brain components associated with vision (Fig. 1)—including *tractus opticus*, *striata*, and *corpus geniculatum laterale* [LGN]—and was strongest between ARDI and the LGN (*p*<0.01). Additionally applying ARDI to brain volumes unveils an association with the visual cortex (*p*<0.1), but only weakly confirms positive associations (Figs 1 & 2b) with areas related to spatial capacities (*hippocampus*, *p=*0.3; *schizocortex, p*=0.4). When spatial areas were aggregated, their correlation with ARDI was moderate (p=0.2) and consistently positive across all five subcomponents (Fig. 2a). Tone and interval had particularly strong associations with the weighted mean of spatial areas (p∼0.1). Subcortical areas involved with motor control were more mixed in comparison with ARDI (Figs 2b & 3), with syllable, transposition, and interval showing the highest correlations (p∼0.2). Basal ganglia (including only *striatum* and *subthalamus*) was surprisingly negatively associated with ARDI whereas *thalamus* and *mesencephalon* (midbrain) were positive (*p*=0.1). If tarsiers are removed from these regressions, the trends are reversed into positive associations. While hormonal areas showed only weak correlations (with interval, p∼0.3), emotional areas had strong positive correlations both interval (p∼0.02), and transposition (p∼0.03) but only a moderate association with ARDI. Other lower brain areas that showed strong associations with areas including motor control (*nucleus basalis*, *p*<0.3 and *vistibularis*, *p*<0.4) as well as those involved in motivation (*habenularis p*=0.01), sleep/wake (*pineale p*<0.001), and arousal (*cortico-basolateralis p*=0.2)--although these were primarily driven by *Tarsier spp*.

**Figure 2.**
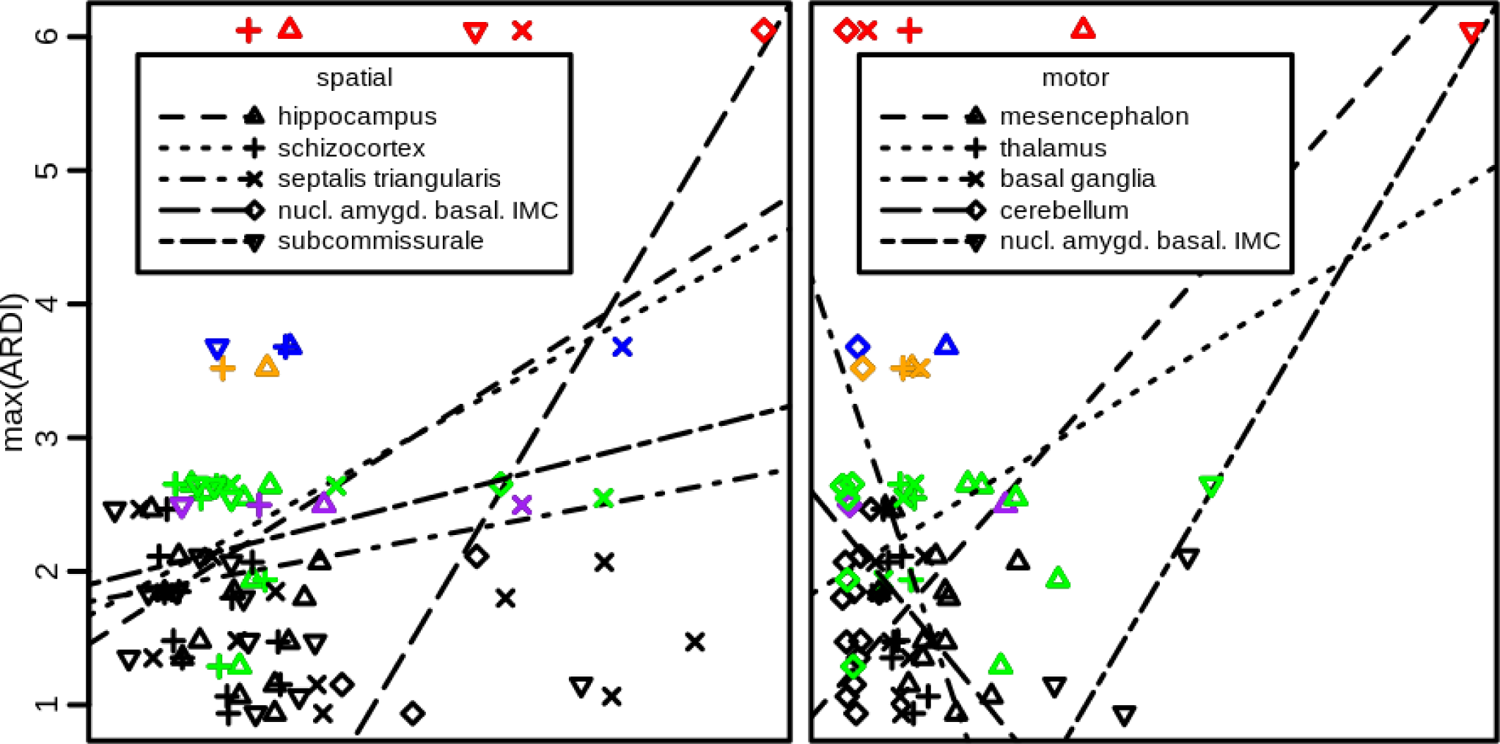
Spatial and motor brain regions associated with ARDI Unaggregated volumetric fractions (per non-neocortical whole brain volumes) from individual brain components (x-axis) plotted against ARDI scores. Linear regression trend lines are individually plotted for each sub-component of motor and spatial areas. With the exception of the basal ganglia and cerebellum, all spatial and motor associations are positive.

## Discussion

A striking, yet unsurprising, primary finding of this work are that primates have diminished neurological capabilities for smell which appears to have been supplanted by spectrally complex vocal communication. This behavioral overhaul, implies possible co-opting of visual areas for “seeing” acoustic gestures, a possible prioritization of spatial-navigational structures (over faculties associated with smell), as well as possible use of vocal signals as indicators for motor control. More tenuously implicated parts correspond to emotive states of motivation, wakefulness, and fear—perhaps regulated in response to and signaled by vocal complexity. Such compensation for arboreally diluted olfactory signaling and inter-substrate leaping, could be related to such ancient factors as target landing, predation avoidance, or some combination thereof, especially in callers.

Socially, I found that higher-cortical paralimbic and medial temporal areas (e.g. schizocortex and insula) use spatiotemporal encoding to link musical stimulii in forming long-term memories for harmonious cohabitation of both trees and forests. These motor control and spatial orientation ideas are integrated under a ‘full cognitive stack’ construct with a discussion on the basal ganglia’s role in motor planning in larger primates, including apes. A chronology of the accrual of individual features underlying musical behavior is explored, with special attention to the development of longer rhythmic and song-like musical production. I conclude by examining the adaptive significance of musicality in [terrestrial] humans with respect to sub-features that are likely in decline (e.g. spectral) versus those possibly under positive selection (e.g. temporal) as well as those being maintained (e.g. syllable).

### Indicators of motor control

The clearest motor-control areas associated with ARDI are the *mesencephalon* (midbrain) and, to a lesser extent, the thalamus (upper *diencephalon*). The midbrain relays sensory-motor signals to the cerebral cortex while the *thalamus* relays movement and motor planning as well as sensory (e.g. vision and hearing) information. It is possible that this increased importance of precise (post-cranial) motor control (e.g. of wrists) co-evolved with expansion of distinctly elaborate acoustic display. Stronger support for (cranial area) motor control is evidenced by greater vocal complexity association with lower-brain areas of *nucleus basalis* and *vestibularis medialis*. The former is known to release the motor-neuron activator acetylcholine, whereas the latter is involved in fine control over head, neck, and eye movements. This evidence supports hand-eye coordination and locomotor signaling origins for our ancestral musicality. The preceding motor-control argument for such a musicality-origins module is nicely complimented by corroborative associations with spontaneous visual-spatial circuitry running conterminously with the diencephalon (e.g. LGN).

### Spatio-social mechanisms across the mate-choice life cycle

Charles Darwin envisioned the evolution of music as a sexually selected trait strongly embedded in the social contexts of “imitation, courtship, and rivalry” (Darwin, 1871). Since then others have proposed various formulations—including pair bonding, group selection, territoriality, and range defense—as inter-related mechanisms driving the evolution of musicality, in myriad species. Here I attempt to bring all of these social mechanisms into a single coherent picture organized around the mate-choice life cycle (Fig. 4) including courtship, copulation, fertilization, parenting, and weaning—from ME to PIANO.

**Figure 3.**
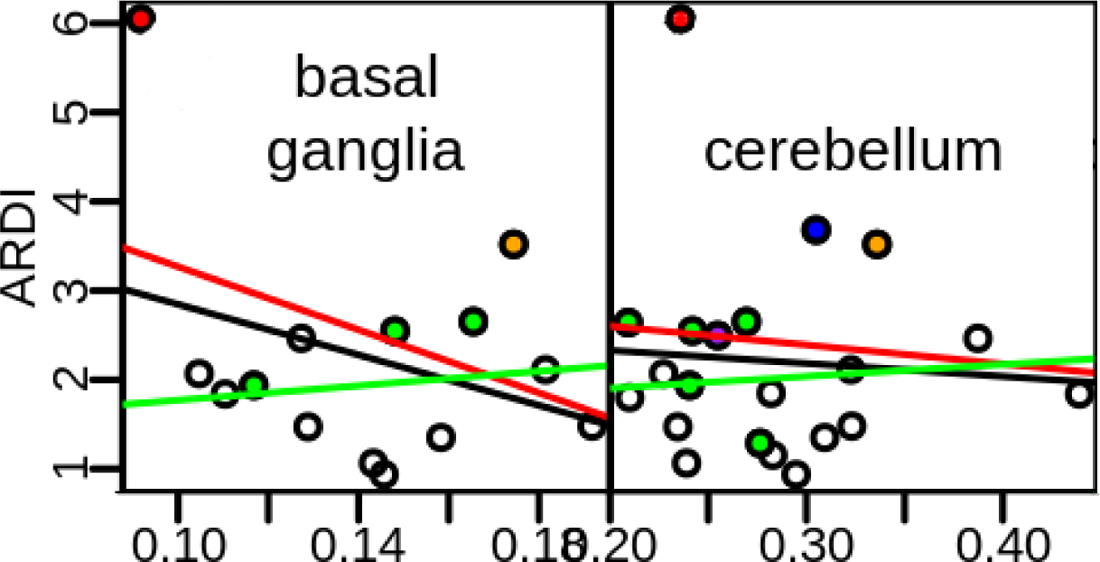
Detailed plots of two contraindicative spatial brain regions. The *Tarsier* outlier drives the negative association of basal ganglia and cerebellum volumetric fractions with ARDI. But *H. lar,* the only gibbon, has high values for both both volumetric fractions and ARDI, indicating motor planning could be higher in larger (e.g. anthropoid) primates than in (e.g. tarsiers). Colored points represent “musical” primates: green=titi, tamarin, and marmoset monkeys, purple=galagos, blue=indri, orange=gibbon, red=tarsier.

**Figure 4.**
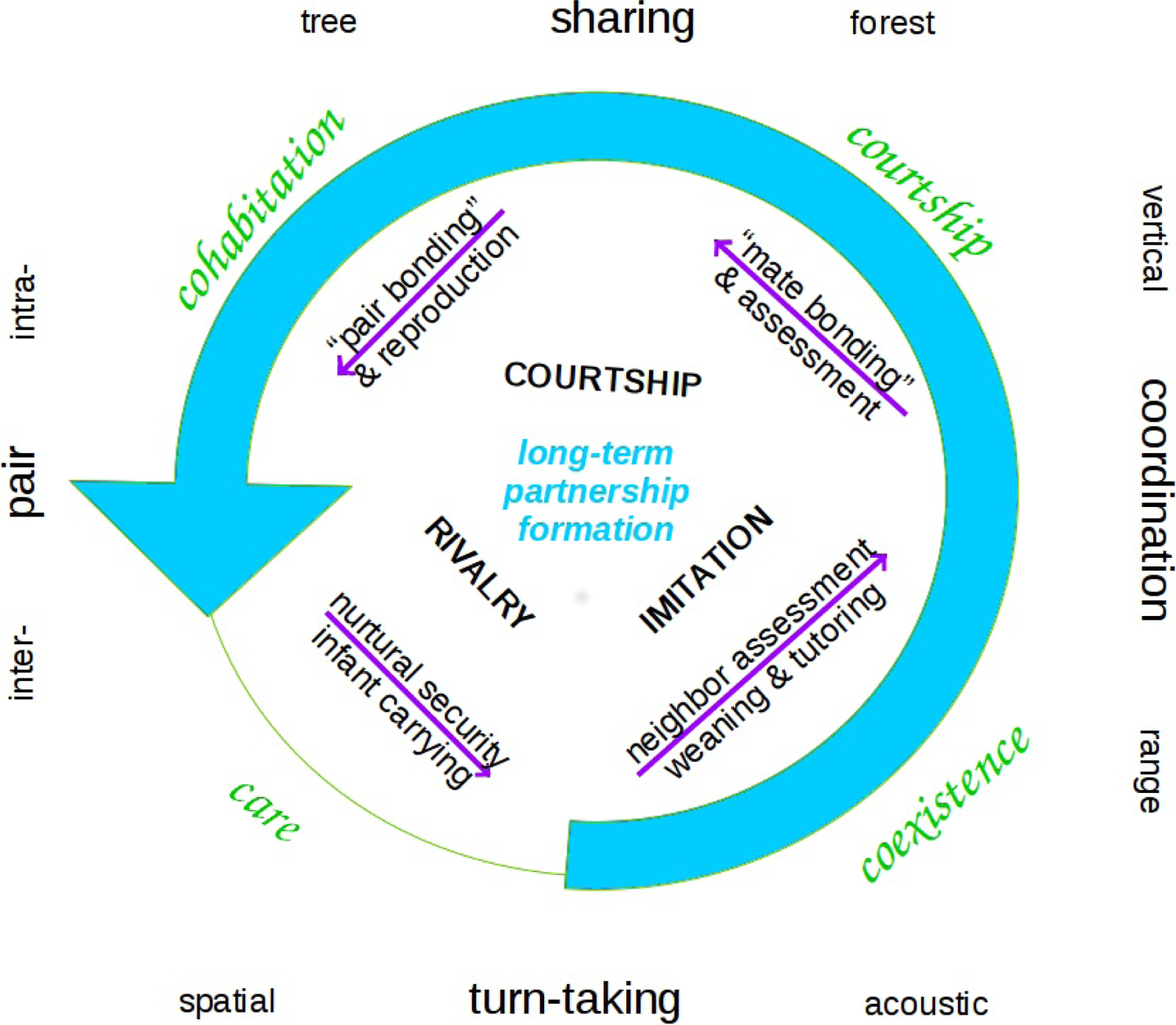
Mate selection, arboreal cohabitation, care, and range spacing. Numerous proposed mechanisms of musical evolution, including Darwin’s original socioemotional contexts, can be organized into a behavioral-ecological mate-choice life-cycle under the four phases of *courtship*, *cohabitation*, *care*, and *coexistence*. While *care* (including ‘nurtural security’ and infant carrying) is excluded from partner formation, it arguably constitutes the end goal of such long-term mutual pairings. The cycle repeats when weaning infants begin learning and imitating the musicality of parents and are overheard by potential mates in neighboring ranges.

### Courtship: remote mate assessment and bond formation

Darwin originally viewed courtship between the sexes to be the original function of the music-like calling in our lineage. Like myself, he preferred to focus on the social dynamics of gibbons as a model organism, from which to rest his ideas on human socially-monogamous courtship.

The association (here in primates) of intervallic vocalizations with hormonal and motor control areas of the brain (e.g. mid-brain and thalamus), may imply that these call features have usurped scent as the primary way breeding pairs communicate in order to discover mutual interest in one another. Naturally, this motor control element of adjacency to highly intervallic calls (Fig. 5), like those of gibbons and tarsiers, reflect fine motor control of the small muscles. As stated in the ME hypothesis above, these vocal motor control signals could indicate functioning muscular control for inter-orbital visual resolution of relatively narrow substrate targets, including branches and tree trunks, for more acrobatic landings. Because both lemurs and gibbons perform daring inter-arboreal airborne feats— where gravity demands split-second visual resolution and skeletal execution of limb landing—effective motor-control signaling using lower brain areas could be a primary function of highly intervallic song.

**Figure 5.**
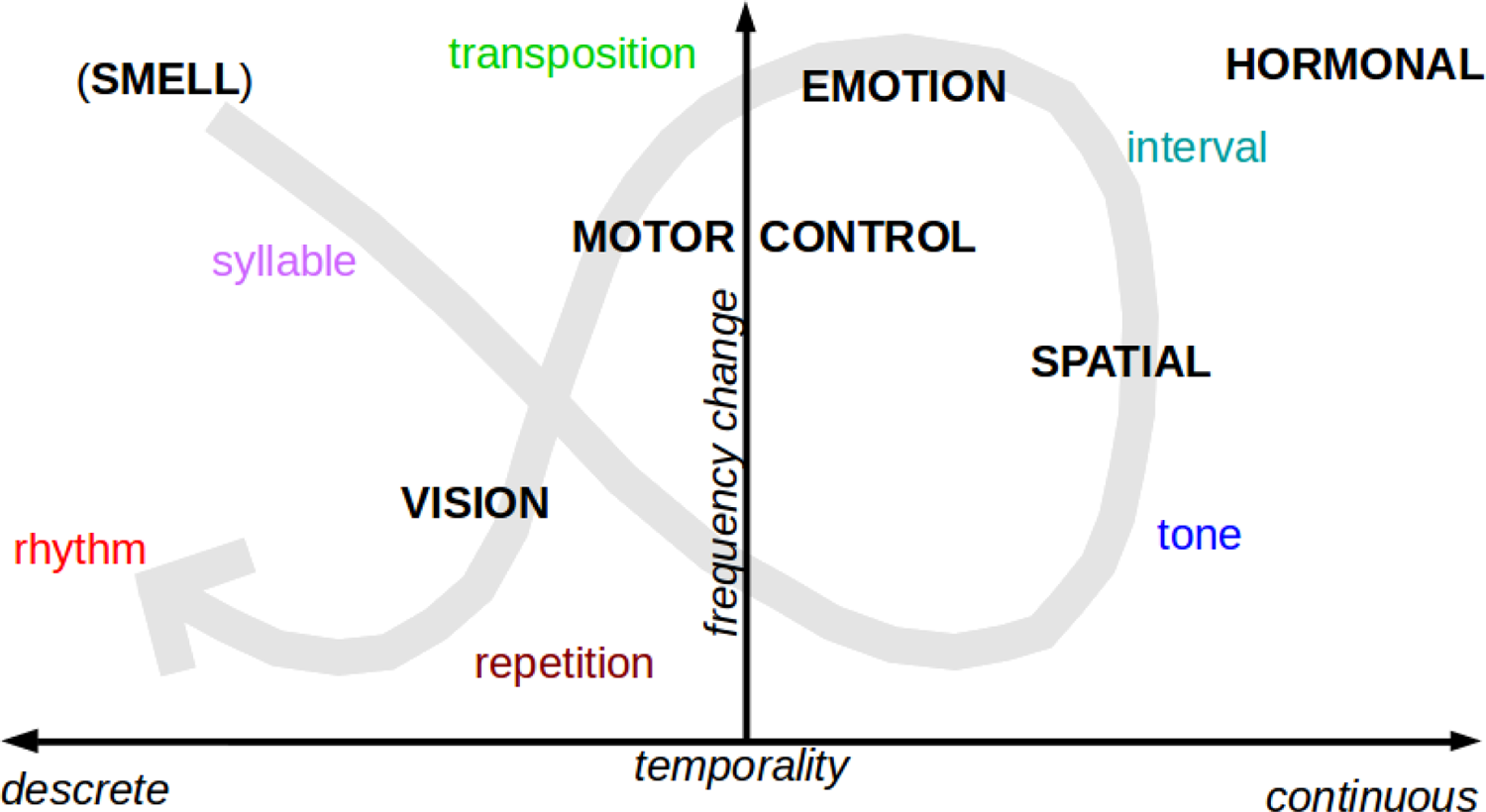
Musical features and brain areas mapped along axes of temporality and frequency dynamics. Musical feature score categories of tone, interval, syllable, repetition, transposition, and rhythm (colored font) and general brain functions (bold all-caps) are mapped in relation to each other and along two types of temporal (reddish) and spectral (bluish) delineations. Temporality (x-axis) distinguishes highly discrete features, like rhythm and syllable, from continuous ones, like tone. Frequency change (y-axis) distinguishes highly intervallic features, including transposition and interval (top), from features that do not modulate in frequency, like repetition (bottom). The evolution of musical signals from the earliest primates to modern humans likely took a path (gray arrow) from reduced smell to a more visually occluded spatial- and tonal-signaling, to intervallic motor signaling, and then ornamental visuomotor control signaling manifested as repetition and rhythm. Parentheses (around ‘SMELL’) indicates a negative correlation.

As discussed in the introductory material on musical production and perception above, indicators of [ME] areas possibly reside in the same general sensory-motor control centers where humans experience pleasurable involuntarily chills, namely just inside the back of neck. These proposed areas possibly include the mid-brain (e.g. PAG, tegmentum, optic tectum, and colliculus) as well as the thalamus.

Socio-ecologically, this mechanism of selecting a sexual partner for their genetically encoded motor-control reflexes, could theoretically, and likely does, happen well before such individuals even come into close contact. Singing (one’s own) and listening to others’ song components could preemptively facilitate a coordination of the timing logistics of sharing acoustic space, remotely, before even sharing a tree or having an up-close encounter. Even frequency based aspects of joint calling could be determined before ever duetting together.

Ultimately, like any k-selected species, longer term signals of future commitment and care are required to transpire between mates as well as short term motor signaling. K-selected species, those with few offspring, invest more in each than r-selected species. Thus potential mates likely factor infant-care and tree-cohabitation considerations, in order to perform precarious joint-arboreal locomotion. Therefore, both ME and PIANO related signaling factor into such long-term selection processes.

### Coordination: locomotor logistics of copulation and bonded cohabitation

While short-term copulations are likely fundamentally rooted in transferring genetic material, even such short encounters require intra-arboreal coordination. Even if parenting and monogamy may not necessarily require constantly sharing or even sleeping in the same tree, copulation by arboreal primates certainly does. While signaling ME to potential genetic mates is important, signaling of PIANO capabilities should be as well. Thus, both working out kinks of (e.g. sometimes even one-armed) arboreal mating [ME] as well as tree cohabitation [PIANO] are arguably both critical social aspects of facilitating any type of pairing. However, while the N in PIANO defaults to *neighbor*, it could also stand for *newlywed*, as my primary focus is on such long-term pairs that spend extended time together in select trees, executing coordinated locomotion, than with any other individuals.

PIANO is also fundamentally an orientational and navigational module which helps couples quickly identify and relocate to one another upon regular daily periods of separation and reunion. Therefore, it ultimately constitutes a form of contact calling ideas on the origins of musicality. However, PIANO is specifically proposed to enable rapid identification of specific spectral features of long-term partners. These features are most directly tied to emotional centers and may even have been co-opted from maternal-infant locational signaling which was only later co-opted to long-term mates (Miani, 2015).

### Care: direct signals to mates realized as parental investment

While every phase of the mate-choice life-cycle leading up to this point has been related to building bonds, the ultimate point of all bonding is arguably to then invest such coordinative efficiency into rearing offspring. That is, in order to turn the caloric payoffs of scouting, nurturing, and even provisioning, into manifest offspring, pairs must put those signaled promises into parental action. Stated using a more classical formulation, there exist adaptive challenges in the logistics not only of mate *selection,* but also mate *cooperation,* in order to successfully bring forth offspring (Zahavi and Zahavi, 1999). This form stands in marked contrast to more recent forms of [credible] signaling, such as from parents who reliably indicate attention towards infant offspring (Mehr and Krasnow, 2017).

Researchers have also investigated different variations on a theme of parental investment related to music evolution. Ellen Dissanayake proposed that [multi-modal (vocal, visual, and gestural)] maternal behavior had direct fitness benefits for improving offspring survival by reinforcing the mother-infant bond (Wallin, B. Merker, and S. Brown, 2000; Dissanayake, 2000). Under more recent views, introduced above, costly forms of parental attention [PA] constituting honest attention can signal PA *to* infants and are indicative of devout infant care and manifested as infant-directed song (Mehr *et al*., 2018). This thesis was recently incorporated into the “credible signaling” framework where parental attention is signaled to helpless infants via simple lullabies between parents and infants (Mehr *et al*., 2020) and where crying (distress, separation, isolation calls) carries information about the timing of investment. While these ideas sound similar to PIANO, my proposal constitutes a signal of promises of future parental investment to potential mates, usually between a male partner to a long-term female partner with whom he has yet to pair. In contrast to conflicting interests between human mothers and infants—a model that fails to recognize the similarly high degree of [arboreal] helplessness of primate infants—the form of PI signal I explore is between potential mates.

This form of PI exposes a different type of negotiation—that *between* sexes—in the form of a possibly collaborative division of labor. Where more ‘gravid’ (or ‘gravis’) infant carrying (predominantly by the female) is juxtaposed with the future nurtural roles assumed by the (presumably more nimble) male, including of food-scouting and range-defense. Although this form of specialization need not necessarily be so sexually asymmetrical, as many male gibbons and callitrichids will carry infants as well as females. But role specialization typically manifests as infants exhibiting a higher degree of attachment to mothers, especially in clades that carry ventrally, and have closer proximity for prolonged nursing. As primates breastfeed for years, in some apes for example, such a reciprocal specialization can manifest with the female carrying and feeding while the male performs scouting duties and defense of the range which his mate and infant occupy.

The signaled promise between pairs is that of PI for nurtural security of the offspring, where both help to convert the calories, afforded to them by their home range, into successful growth of their progeny. Nurtural security gets at the heart of evolution because it focuses on turning genes and food into embodied offspring. Because dietary factors affecting mate choice are highly confounded with other measures (e.g. size, health), it is often instead assessed analytically via more direct measures of home range and care.

In other musically singing animals like birds, gametes (here, eggs) are exceedingly larger in the female and she is therefore undually burdened by extra weight for flying, and therefore that much more vulnerable to predators. It remains unclear how this logic might affect larger cetaceans, except perhaps in that they may also face a similar form of obstetrical dilemma when carrying fetuses (or similar slowness as a pod) that could interfere with efficient swimming. Thus many musical species could have evolved courtship signaling between pairs in order to evaluate promises of effective cooperative role specialization in prolonged parental care contexts.

### Coexistence: inter-pair re-orientive spacing upon pair relocation

Coming full circle back to weaning and independent feeding by offspring, established pairs likely also begin to explore their own range and the boundaries with others’. Such calls began to assimilate by virtue of physical attachment to their mother may begin to manifest as their own ranging calls. These short (not necessarily repeated) melodic, and initially just intra-group directed, calls may function in a spatial orientating [PIANO] capacity.

With locomotor independence, comes visuomotor competence in resolving branch shapes for themselves. Thus, even though calls serve in a PIANO capacity for orientating towards conspecifics, experimentation with longer more repetitive calls may also signal mid-brain and thalamic function used in binocular vision for branch-shape resolution and limb-landing.

But as a k-selected long-term family unit, these sub-adults may still be dependent upon their parents for physical protection and knowledge-based seasonal foraging. In visually (arboreally) occluded habitats, not only spectral precision but spatial orientation is required. Both the archicortical/limbic hippocampus, allocortical/paralimbic entorhinal areas are involved in processing the full range of auditory stimuli, any initial attempts at auditory grouping, and potentially also spectral binding of acoustic elements. Researchers have demonstrated negative associations between dissonance and hippocampal regions, which could possibly be interpreted to undermine any affiliative consequences implicit to PIANO. Noisy, dissonant, short staccato notes are the key parameters that characterize alarm calls (Snowdon 2001), whereas affiliative contact calls tend to be more harmonic (Fischer 2000)

But it’s possible that these more dissonant alarm calls could have been counter-intuitively co-opted to stimulate range-spacing and reproductive *suppression* in neighbors that have been perceived by the callers as encroaching upon their range. Use of harsher and more dissonant alarm and aggression calls may stimulate (such neighbors’) nervous systems at the (sub-)thalamic level (Blood and Zatorre, 2001) and possibly into reflexive (ME) response for fight or flight reaction. The data presented here further support highly intervallic calls possibly replacing smell as a primary means of stimulating sub-cortical brain areas (e.g. amygdala) associated with the fear response. Such mildly alarming acoustic stimulation could also possibly act to suppress reproduction and thereby facilitate inter-range buffering. That is, dissonant or piercingly intervallic calls could directly impart to neighbors signals to relocate or otherwise suppress reproductive impacts on overlapping habitat. While highly intervallic calls are not dissonant in human music, in other animals they may be considered as both dissonant and yet paradoxically also cooperatively affiliative, in this proposed form of range-spacing coexistence.

In contrast to the harsh crescendoing frequency sweeps of territorial gibbon great calls, adult quaver phrases of this ape family are considerably more assuaging. These quaver phrases may be understood as more affiliative, and may serve as a more acquiescent adult (PIANO-esque) signal—possibly, for example, when and where to relocate or cross gaps in the canopy. This consideration reflects a shift from slightly lower-brain hippocampus (and amygdala) level processing (e.g. stimulus binding) of harsher territorial signals to those that are higher and more distant from the basal forebrain. The insula may be recruited to form a more gestaltically coherent environmental scene for planning more elaborate, unusual, and risky locomotor maneuvers. Also, while still medial, the insula is lateral of much of the limbic and paralimbic areas involved in more simplistic processing of stimulus. The insula is responsible for more socially-informed emotional processing like compassion, empathy, and interpersonal self-awareness. This functionality corroborates the typically affiliative nature of most music-like calls and higher brain appreciation and awareness of others, as well as possible suppression of lower-brain reproductive urges.

A final piece of the coexistence theme is that of imitation, or mimicry, of the calls of neighbors. In birds, songs tend to function as signals (or “threats”) of readiness or willingness to defend their territory. Mimicry of a neighbor’s song addresses the threats of that particular neighbor. It also gives the singers a chance to show off their age, experience, learning, and memory (in a very directed fashion) in their reply to a specific neighbor (Zahavi and Zahavi, 1999).

Charles Darwin thought of music as the supreme mystery of the science of man and the most mysterious trait with which we are endowed. He also thought of music as the best example of mate choice playing a role in a human behavioral trait (Darwin, 1871). Geoffrey Miller has been Darwin’s most vociferous supporter (on this topic) in the last hundred years (Miller, 2000). He has championed Darwin’s view on sexual selection through mate choice for brains as indicators [of fitness] as a way to preempt natural selection (Miller, 2001). He has viewed sexual courtship as the principal biological function of music, dividing mate choice into either indicators or aesthetic display (Miller 2000). Here have I have proposed a third way, via PIANO’s spatial reciprocity in long term parental investment through care as nurtural security, or infant carrying, or both.

### Paralimbic / Insular Acoustic Neighbor Orientation versus ASA

There has been a respectable research effort levied against the theoretical framework of *auditory scene analysis* [ASA] (Bregman, 1990; Trainor, 2015). But very little of this work specifically addresses spatial aptitude aspects of cognition in the signal sender. ASA instead focuses on distillation of contiguous shapes across topographically equivalent representations of cognition (in the signal receiver). I suggest a refocusing toward neural representations (of *both* the signaler and receiver), specifically those interconnected to the (para)limbic regions of the brain.

I propose a renewed and expanded reconsideration of ASA in both how and why the auditory scene is connected to specific cognitive systems. I propose that an ecologically relevant capacity for orientation within an organism’s acoustic neighborhood—one comprised of uniquely (and perhaps musically) identifiable auditory stimulii—could plausibly run conterminously with the paralimbic system. It is further possible that such ancient, and *olfactory* contiguous, cortical structures were repurposed to encode pertinent *acoustic* landmarks for orientation, especially with regards to conspecifics. This is in spite of the fact that evidence presented here—for any such putative exaptation of olfaction-adjacent regions towards acoustic processing—is still quite limited.

### Ancient primate homology and the multifactorial evolution of music

Recent studies of the brain provide evidence—in the form of circadian activity period, respiration, and even brain waves—that rhythmicity is ancient in animals. Some have further suggested that processing of rhythm has deeply ancient underpinnings (Honing, 2019). But, while my own investigations into individual musical features confirm monotonic rhythm is pervasive in primates (Schruth, 2022), higher order rhythmic complexity has increased only very recently in larger more terrestrial forms. In other work, I have demonstrated how a minimalistic definition of human musicality (e.g. as ARDI) is all-inclusively derivable across all human cultures. This (rhythm and tone agnostic) definition also applies across deep-clades to the origins of the primate order itself, with spectral complexity (e.g. melody) as the default state for primates. Presumably, spectral features replaced chemical signaling in crucial social communications, such as those for mating. Using the neural correlations underlying Figure 1, I outline a rough chronology within the organizing context of both temporality and spectral dynamics (Fig. 5) in the sub-sections in remainder of the discussion.

### From olfactory signals to discretized acoustic referents

The observed trade-off between sense of smell—diminished olfactory and piriform areas—with other areas observed here (Fig. 1) is unsurprising. Because complex auditory and visual processing primarily localize to the temporal and occipital regions of the cerebral cortex, a conversely decreased correspondence in brain parts for reduced olfaction is expected. A more striking observation, however, suggests primates and their predecessors may have adjusted allocation in lower cortical areas, that were originally evolved in ancient piscine ancestors epochs earlier, toward more auditory-based spatial orientating. As mentioned above, ancient cortical areas for smell seem to have been efficiently reduced and out-competed for adjacent space. Also plausible is the idea of overlapping functionality, or shared contiguity, with regions of both higher cortical areas involved in processing of auditory input as well as lower cortical areas involved in spatial processing, perhaps also by recruiting hippocampal regions. These two sensory systems reside on opposite extremes of the well interconnected hippocampal gyrus—with the (lower) limbic areas primarily processing chemical stimuli and the higher temporal regions primarily processing auditory inputs.

### Emotional cries, wails, and moans: weaning, attention, and predation

The results of the regression of brain volume fractions upon complex calling most strongly implicates diencephalic-contiguous areas including the midbrain and forebrain. Due to the disproportionately extreme values for *Tarsier spp.*, these parts—including the *pineale* (sleep/wake), *amygdala* (arousal/fear), and *hebernularis* (motivation/addiction)—exhibited strong associations with musical calling. The strong associations of the pineal gland and habernularis with musical complexity suggests PIANO could benefit from expanded consideration to lower brain areas involved in wakefulness and arousal. But strong associations of vocal complexity with wakefulness are not surprising, especially considering a reduction in ability for remotely sensing the emotive states among arboreally dispersed and perpetually repositioning (e.g. non-nesting) conspecifics. Each of these could factor into an broader model including both sensory tradeoffs and predation-avoidance strategies (Schruth and Jordania, 2020; Schruth, 2021).

As a final consideration of inter-arboreal pressures and predation avoidance driving arboreality and eventually music-like calling, I also address the puzzle of culogos. Culogos are tropical arboreal inter-tree gliders who, along with tree-shrews, are close contenders for being the most closely related taxa to the primate order. It is natural to suspect, given the logic of both ME and PIANO outlined above, that culogos should likewise have developed musical calling, given their inevitable gliding-driven physical separation and high-velocity limb landing. However, I would argue that several key differences have stifled such an evolution towards musicality in their lineage, including claws instead of nails, reduced infant carrying, their gliding anatomy, and their infrequency of inter-tree leaps. Primates have nails which presumably make infant carrying more comfortable. Cologos, by contrast, have slightly higher parity as afforded by leaving their infants in nests instead of carrying them. Furthermore, cologos only glide for a very small percentage of their overall locomotion, in comparison to the more than fifty percent for leaping by lemurs. Lastly, gliding has major glide-landing based advantages over primate leap-landing. First, cologos’ targets (e.g. tree trunks) are broader and therefore presumably easier to land on. Secondly, cologos’ landing forces may be more gentle, considering their parachute-style landing, when compared to wingless primates. Thus, the lack of an evolved musical signaling capacity in cologos could be explained by lower-stakes landing implications of not only their life-history but anatomy.

### Tradeoff for sense of vision

A similar co-opting of visual cortical areas by cognitive processing of music-like gestures, are arguably evidenced here. It is possible that nocturnal and leaping near stem-primates repurposed the *striata* (Fig. 2), a cortical area normally used for processing optical inputs, for visualizing auditory input instead. Something analogous is known to occur with elite, but visually blind, musicians (e.g. piano players)—who recruit the occipital lobe in facilitating musical performance (Ross, Olson and Gore, 2003). The similar phenomenon of mental representation of exogenous spectral contiguities to internal spatial cortical maps has been covered in *auditory scene analysis* [ASA]. However, much of ASA is typically concerned with tracking of continuous streams of auditory sensory input. In contrast, I explore matching of discrete acoustic gestures for limb landing (e.g. ME) and mapping of these gestures into ecologically relevant spatial coordinates (e.g. PIANO).

### Visuospatial signals

The strongest evidence of a correlation between any of these vocal complexity scores and visual brain structures (Fig. 2) lies, interestingly, in the LGN. The LGN obtains information directly from retinal ganglion cells. It also has strong feedback connections with the visual striata. Thus a large LGN indicates exceptional prioritization of visual input and a need for instantaneous spatiotemporal calculations, perhaps for rapid motor reflexes. Further, the LGN also plays an important role in the integration of directives for visual attention with other sensory input, such as sound cues. Output signals from the LGN determine the spatial dimensions of the raw visual inputs from anatomically corresponding points along both retina. This information is used as feedback for visual focus on objects of interest in the external environment. Thus, the LGN could serve in locating conspecifics who might be separated in arboreal, visually-occluded, environs where transposition-rich musical calls would propagate most efficiently.

### Terrestrial Ecology

Spectral attributes of acoustic output seem to decline for most terrestrial species. But syllables may be maintained via neutral or balancing selection. In humans, who have noses that rise meters above the (relatively odoriferous) earth, such selection may have even been positive. But other spectral qualities are likely undergoing decline as they have in other terrestrial primates and birds. A few exceptional possibilities causing (spatio-) motor emplacement [ME] to persist in humans include throwing as part of ballistic hunting in humans. Features of PIANO also could have applied to more nomadic hunter-gatherers or pastoralists who also needed to orient to repositioning of nearby tribes—ever-shifting acoustic landmarks on the nomadic ancestral plain. A more postural application of ME could be hammering for stone toolmaking (Schruth, *et al*., 2020). Even more intricate tool crafting, such as those that don’t fossilize or last millions of years, could also have comprised the underlying trait bolstering such signaling via musicality. These other less ballistic forms may have entailed more complex motor plans with numerous lengthy steps and could have corresponded to longer more hierarchically elaborate songs. This scenario may then Implicate a final missing piece in a full stack of vertically connected brain components—including cerebellum and basal ganglia—which lie intermediate to areas involved in ME (inferior/medial) and PIANO (superior/lateral).

Yet evidence suggests that only temporal qualities should increase, and then for max, not mean levels of patterning. Thus even such temporal acoustic signals in humans may have been ornamental for sexual signals rather than for routine egalitarian ones. If larger visual areas are primarily associated with repetition, for example, and my proposal concerning interorbital pattern-matching (as part of ME) is correct—occasional rapid climbing could have been the primary target of (indirect) signal selection. This scenario is most coherent if early *Homo* used trees only for sleeping and protection during the night. Even later hominins would likely still occasionally climb trees to gather fruit or honey. This preservation of repetitive acoustics as a sexually selected ornament in males would be compatible with provisioning behavior by males during high calorie demands of gestation and lactation. Furthermore, assuming a more gibbon- or bonobo-like, free and egalitarian mating system, an increasing sexual dimorphism in our lineage could explain the carryover into the modest imbalance of singer sex-ratios in modern human music.

### Sexual selection and temporal display

There is a conspicuous peculiarity regarding the popularity and fervor us modern Westernized humans have for our music, particularly because our practice is so commonly decoupled from any obvious survival value. Even the astonishing genius of Charles Darwin struggled with the problem of music in humans and accordingly allocated decades of additional labor to tackle such curious emotional traits, inventing a new form of (sexual) selection, en route. That its ultimate function is so mystifying is a major clue that he may have been on the right track by looking at the differences in mating goals of sexually dimorphic species. I have argued above that musicality arose in more committed partnerships: in k-selected species, for the mutualistic goals of rearing offspring in arboreally precarious habitats. But I have also shared evidence that terrrestriality tends to lessen the very spectral qualities that are signaled by an increasingly irrelevant form of locomotion. Thus, while musicality may have initially evolved via sexual choice or mate selection for monogamous arborealists, terrestrial descendant appear to have only weakened any pre-existing manifestations. I have proposed that formerly arboreal and now terrestrial species, like ours, are experiencing a split between vestigial spectral qualities of their musicality.

However, I have also reported some features of temporal patterning that have increased. For example, the average of the most rhythmic calls in terrestrial species is more rhythmic than the average of the most rhythmic calls in arboreal species (Schruth, 2022). Thus, it appears that temporal features of terrestrial primates are becoming more ornamental and perhaps therefore also dimorphic and truly sexually selected. A case in point is the display of a chest beating gorilla who is habitually terrestrial and may only very occasionally need to use abstract pattern matching (for ME) to rapidly climb a tree, perhaps in rare circumstances when not the largest animal around. This form of terrestrial signal, where upper and lower body motor functions are decoupled, could be the norm for signaling full functionality in such semi-vestigial limbs (Schruth, 2022). Similar dimorphism appears to have appeared in more rhythmic displays of humans (e.g. dancing).

A higher degree of rhythmic patterning of vocalizations has been observed to result in greater reproductive success in birds. Longer more rhythmically-precise singing may signal superior status in being unperturbed by predators in spite of giving away location. Uniformity of trilling may likewise signal confidence as only those individuals who can afford to concentrate full attention on their calling are able to reliably make a pattern (Zahavi and Zahavi, 1999). Thus rhythm, like melodic matching, may have similar signaling mechanisms, here illustrated via more costly rather than indicative, for branch landing.

### Group factors on motive emplacement and entrainment

This paper began by advancing the weak assertion that motive emplacement was the neuro-modular substrate acted upon by indirect selection for genes by the opposite sex. And while I continue to argue for higher brain processing of such signals in a way that facilitates higher-order pro-sociality (e.g. child-rearing and spacing), here I return, full circle, to motive emplacement, but now in a larger group context. The primary way in which entrainment could provide value is through the context of rhythmic group performance. But instead of placing limbs on branches, we now grasp each other. Dancing (by pairs) has arguably become a (loco)motor-visual display using semi-vestigial limbs accompanying temporally dominating musical-auditory signals. These signals possibly evolved to demonstrate that such limbs still can, in-fact, function, for other occasional secondary (non-locomotor) uses.

Despite the recent finding that rhythmicity tends to decrease as group sizes grow larger across primate species (Schruth, 2022), there is something alluringly inviting about a (highly conspicuous) instrumental rhythmic performance. Other researchers have noted that metrical rhythm is easy for an outsider to join (Savage *et al*., 2021). I have recently noted that there even appears to be more room for additional performers beyond the typical small-family spectral-performance of gibbons, to something on the order of six individuals, in more rhythmically-inclined species (Schruth 2022). There is a strong relationship between terrestriality and group size, and for our progenitors to survive terrestrially, they needed to learn how to work together in larger groups: for example to team up on more ferocious terrestrial carnivores, for scavenging, or hunting, or both.

Thus musicality seems to run the full gamut of associations from those involving infant care, to mate bonding, to intra- and inter-group signaling. I thereby concur with efforts to develop a broad and inclusive theory where signaling and bonding theories can coexist under one framework (Savage *et al*., 2021). Others have argued that this spectrum is actually U-shaped from mother-infant to pair to group (Lehmann, Welker and Schiefenhövel, 2009), more akin to what I have attempted to do above from a mostly signaling perspective. And yet, concluding with the curious case of human music, this paper comes chronologically full-circle back to short-term sexually-selective signaling for copulation opportunities presented by a visuomotor flavor of the ME module, described above. Rhythmic patterning has turned out to be not only the most ancient (monotonic), but paradoxically also the most modern (complex). And while there were not strong correlations for monotonic rhythm, others have noted an important role of the cerebellum and embodied motor areas.

### Motor planning, basal ganglia, and the full cognitive stack of music

Above I argued that MEE may have been important for movement in precarious proximity, such as in acrobatic tree-sharing by gibbons or other aerial-prone primates. But its not entirely clear why MEE might be important in humans, except for the occasionally intense bout of hand-to-hand fighting. An answer could come in the form of complex motor planning and execution. While vocal production learning is largely absent in primates (Janik and Slater, 1997; Fitch, 2015), sequencing of song elements (Petkov and Wilson, 2012) may be seen in certain species. Primates are notorious for being unable to use their lips and tongues in a way that comes close to what humans are capable of when using language. But apes are extremely proficient at communicating using their hands. This has led some researches to even go as far as defining musicality as “deliberately controlled and metrically organized gesture” (Morley, 2014). There exists an extraordinary similarity between music and language (Merker, Morley and Zuidema, 2015) with corresponding requisite motor sequencing (Brown, Martinez and Parsons, 2004) and the maintenance selection on syllable diversity— both possibly translating into lexical processing areas of the temporal lobe (e.g. Broca’s & Heschl’s areas). This line of thinking, however, may require a punctuated leap in abilities from the motor-planning of limbs (in apes) to that of the vocal apparatus (in humans). Furthermore, the evidence for flexibility, and thereby planning capacities, for vocal production in primates is mixed at best. Some plasticity has been demonstrated in spectral features of calls in marmosets and tamarins (Weiss, Hotchkin and Parks, 2014) as well as for loudness and duration in macaques (Bryant, 2013). Gibbons have a high degree of heritability of a fixed syntax to their calls (Geissmann, 2000; Fitch, 2015), although the spacing, timing, amplitude and spectral qualities are obviously refined between infancy and adulthood.

Duetting in primates is a rare form of acoustic display that requires close coordination with a partner. And while others have investigated (*temporal*) social syntax turn-taking in anthropoids (Snowdon, 1990; Falk, 2004), few have explored *spectral* aspects of coordination. Harmony is mentioned in the context of musical (conversational) turn-taking (Dissanayake, 2000) and pitch-matching (Cowley, 1998). In contrast, I propose that (non-overlapping) *spectral* harmony has roots in signaling maturity of [arboreal] space-sharing cognition for precarious joint-locomotor activity—especially in gibbons, tarsiers, and some lemurs. Such cases of spectral interleaving, however, rarely extend into longer complex sequences.

The basal ganglia [BG], an area thought to be central for complex motor planing, was negatively correlated with the ARDI musicality measure. A salvageability of flexible planning capacities is possible, however, if we can assume (more predictably vertical trunk-landing) tarsiers represent an exceptional outlier. If the BG facilitates both complex locomotor planning—including convoluted (horizontal) branch landing in gibbons or complex multi-person (e.g. swing) dancing in humans—it may represent a recently completed ‘full stack’ of motor control underlying musical signaling. Thus human music may represent a full cognitive assemblage of motor control modules through many levels, cortices, and systems.

### Adaptive significance to modern societies and civilizations

Music has been described as a non-selected by-product of biased predispositions for other evolved capacities. Steven Pinker famously called music “Auditory cheesecake.” Indeed, music could prove to merely be a culturally created yet highly evocative system of sounds designed to exploit unconscious and biologically evolved perceptual response biases to distinguish the affiliative from the threatening (Bryant, 2013). Just because certain display traits may have originated via non-selective mating preference, however, doesn’t mean they have to remain non-selective (Searcy, 1992). Recently there has been a call to move beyond the idea of by-products as final evolutionary dead ends, towards thinking about exaptation instead (Savage *et al*., 2021). Rather than thinking of music exploiting auditory scene analysis modules, we can invoke culture as salvaging this perceptual apparatus as exaptation (Trainor, 2015, p. 2). In fact, the entire history of musical feature accrual could be re-construed in this way. Music is composed of components that correspond to different adaptive pressures over evolutionary time (Honing *et al*., 2015; Trainor, 2015).

On the opposite extreme, some have claimed that human music is entirely unique. Despite claims that animal musicality and singing has no homology or analogy with human forms (Hauser and McDermott, 2003; Savage *et al*., 2021), the data suggest otherwise. I have shown elsewhere (Schruth, 2020) that musicality (as measured via ARDI) has a strong evolutionary signal (lambda∼0.8) evidencing a deep homology in primates. Further, while our singing is likely not homologous with birds, and constitutes a more rare form in primates, humans do share substantial homology with gibbons. Gibbons happen to have the lowest average of this taxonomic family’s species most rhythmic calls (Schruth, 2022). But unlike increasingly vestigial melody, terrestriality seems to have increased selection for ornamental rhythm in humans. This combined with complex motor planning required for dancing and language will likely show that vocal rhythmic display constitutes a tirelessly fruitful area of study.

## Conclusion

In conclusion, music-like signaling appears to have evolved with fundamental visual, motor, emotive, and spatial-orientating structures of the mid-brain, allo-archi- and paralimbic cortical areas. Associations of temporally aesthetic features with visuospatial areas confirm auditory grouping predictions of both PIANO and ME. These further indicate possible ornamental selection for vision-related function associated with ME in terrestrial primates who practice only semi-vestigial use of forelimbs. Spectral feature associations with spatial areas, notably, may be more relevant in visually occluded environments. Further, these acoustic features have strong phylogenetic signals and their cognitive associations coincide in deep-clade primate lineages such as tarsiers.

The paralimbic and archicortical areas, in particular, seem well situated between the most ancient sense of smell and the more recently evolved sense hearing— both highly integral in localization of the moving landmarks of conspecifics and predators alike. Intervallic associations with motor areas, on the other hand, suggest musical frequency changes in calls could serve as indirect signals for fine muscle control in (also formerly) arboreally acrobatic species. It is possible that cognition for in-motion binocular visual resolution of branch shapes required to accurately land high-speed locomotor bouts—may have also occasionally been co-opted for auditory signal assessment when stationary. The comparative processing of auditory-spectral gestures was likely not only used for identification and localization, but could have supplementally been recruited for assessment of display quality. Such a receiver-side function of display quality assessment, rather than a strictly orientational capacity, is evidenced by the higher correlation of musical calling with orbital-motor (LGN) over paralimbic regions, though both associate positively.

Strong emotional associations with (intervallic) spectral features suggest musical calls may have early maternal-infant locational roots. But these signals may have been co-opted by adults to signal reliable motor control capacities to long-term mates. The model presented here further suggests that (arboreal) signal receivers use spectral qualities of possibly direct signals of nurtural security to assess quality of long-term mates. It is also possible that a harmonious inter-family coexistence driven by range spacing is enforced via hormonal-reproductive suppression via these very same types of (e.g. highly intervallic) calls.

Rhythmic complexity has independently evolved in several disparate terrestrial primates, but in humans it has coincided with exceptionally longer songs and linguistic sequences. Thus while rhythmic aspects may currently persist in humans as indirect opportunistic indicators of occasional motor-control capacities, more fully featured musical complexity (e.g. harmony) appears to have evolved in longer-term arboreal mate-choice consistency contexts. Only through a life-time and life-history holistic perspective, considering the complete mate-choice cycle, can we understand both the indirect and direct selection effects driving musical courtship signaling, for both genetics and nurture of viable offspring.

## Acknowledgements

I thank Rob, Aditya, Sarah, and Jeannie, and Tiffany for scoring spectrograms. I also thank my former collaborators Chris Templeton, Darryl Holman, and Eric Smith for their ongoing contributions to related projects. I also thank Eliot Brenowitz and Joseph Sisneiros for granting me permission to audit their neuroetholgy course. I also thank Ellen Dissanayake for editorial assistance.

## Statement of Ethics

No live animals were used in the compilation of our data.

## Data Availability Statement

The data is available on-line at http://osf.io/bvsfz

